# Liking and left amygdala activity during food versus non-food processing are modulated by emotional context

**DOI:** 10.1101/707844

**Authors:** Isabel García-García, Jana Kube, Filip Morys, Anne Schrimpf, Ahmad S. Kanaan, Michael Gaebler, Arno Villringer, Alain Dagher, Annette Horstmann, Jane Neumann

## Abstract

Emotions can influence our eating behaviors. Facing an acute stressor or being in a positive mood are examples of situations that tend to modify appetite. However, the question of how the brain integrates these emotion-related changes in food processing remains elusive. Here we designed an emotional priming fMRI task to test if amygdala activity during food pictures differs depending on the emotional context. Fifty-eight female participants completed a novel emotional priming task, in which emotional images of negative, neutral or positive situations were followed by pictures of either foods or objects. After priming in each trial, participants rated how much they liked the shown foods or objects. We analyzed how brain activity during the contrast “foods > objects” changed according to the emotional context – in the whole brain and in the amygdala. We also examined the potential effect of adiposity (i.e., waist circumference). We observed a higher difference between liking scores for foods and objects after positive priming than after neutral priming. In the left amygdala, activity in the contrast “foods > objects” was higher after neutral priming relative to negative priming. Waist circumference was not significantly related to this emotional priming effect on food processing. Our results suggest that emotional context alters food and non-food perception, both in terms of liking scores and with regards to engagement of the left amygdala. Moreover, our findings indicate that emotional context has an impact on the salience advantage of food, possibly affecting eating behavior.

## Introduction

Emotions can influence eating behaviors (Geliebter and Aversa, 2003). Specifically, it is well established that *negative* emotions can disturb our food perception and appetite (Evers et al., 2010; Macht, 2008). These effects of negative emotions on eating behavior are nevertheless largely heterogeneous, where around 30% of the population experience increases in appetite, 48% decreases in appetite and food consumption, and the rest is not changing their eating motivation (Macht, 2008). Positive emotions have been less often studied but suggested to influence food intake as well. When individuals are in a positive compared to a neutral mood state, they tend to rate foods as being more pleasant (Bongers et al., 2013; Greimel et al., 2006) and to increase the amount of food that they eat (Evers et al., 2013; Paquet et al., 2003).

Body weight, and particularly obesity, is a factor that might influence the effects of emotional state on eating behavior. Studies have reported that psychosocial events associated with negative emotional states, such as work stress, predict weight gain in participants with a high BMI (Block et al., 2009; Fujishiro et al., 2015; Kivimäki et al., 2006). In lean participants, conversely, psychosocial stress seems to be linked with weight loss or no weight changes (Block et al., 2009; Fujishiro et al., 2015; Kivimäki et al., 2006). Individuals with high BMI also report more frequent use of eating and drinking as a stress coping strategy (Laitinen et al., 2002). These results suggest that emotions influence eating behavior differently in individuals with obesity compared to lean individuals.

In the brain, the amygdala is one of the regions in which food and emotional processing might interact. A meta-analysis of fMRI studies found a robust engagement of the bilateral amygdala in response to negative emotional stimuli (García-García et al., 2016). This brain region is frequently selected as Region of Interest (ROI) in emotional priming studies (Pichon et al., 2012). The amygdala has also been widely implicated in the processing of food and other rewarding stimuli (Sescousse et al., 2013; Tang et al., 2012; van der Laan et al., 2011). Moreover, altered amygdala activity might be clinically-relevant in this context, since both individuals with obesity and patients with substance use disorders were shown to exhibit *increased* activity of this region in response to food and drug stimuli (García-García et al., 2014).

The amygdala exhibits widespread connections with the rest of the brain. Its basolateral subdivision is bilaterally connected with the hippocampus, medial prefrontal cortex and orbitofrontal cortex, and these connections seem to sustain habit-based behavior, cognitive control and reward processing, respectively (Janak and Tye, 2015). The central nucleus of the amygdala projects efferent signals to the ventral tegmental area and to the substantia nigra pars compacta, possibly modulating the computation of reward signals (Watabe-Uchida et al., 2012). Researchers in the neurobiology of emotion and eating behavior have also emphasized the importance of the interactions between the amygdala and the ventromedial prefrontal cortex. This circuit is critically involved in the processing of subjective value, emotion regulation and fear extinction (Clithero and Rangel, 2013; Phelps et al., 2004; Seo et al., 2016). Studies have shown that animals with lesions in the amygdala ventromedial prefrontal pathways show altered decision making (Baxter et al., 2000; Fiuzat et al., 2017; St. Onge et al., 2012).

In sum, the amygdala has been implicated in the processing of both emotional stimuli and food, and it is widely connected with other reward and salience areas. This makes the amygdala a plausible structure engaged in the modulation of food processing according to the emotional context. However, to date there is no study directly addressing the role of the amygdala in this modulation.

In the present study we examine how emotional stimuli modify the subsequent processing of food cues in the brain. We designed an *fMRI emotional priming task* where we displayed sets of food and non-food (i.e., objects) pictures right after the presentation of emotional stimuli (negative, neutral, and positive). The first step was to test whether the emotional images were able to elicit a pattern of brain activity that is in line with the existing literature. Second, we examined the contrast “foods > objects” across the different emotional conditions. We hypothesized that both liking scores and activity in the amygdala in response to food stimuli would vary depending on the emotional context. We also hypothesized that body weight status, measured with waist circumference, would have an effect on the fMRI signal and liking rates.

## Method

### Participants

The final sample consisted on 58 participants with their body mass index (BMI) ranging from 17.67 to 46.83 kg/m^2^. We recruited participants from the volunteer pool of the Max Planck Institute for Human Cognitive and Brain Sciences (Leipzig, Germany). We only included women in the study to reduce potential sex-related heterogeneity in the responses to emotional stimuli and food stimuli. The study was performed in compliance with the Declaration of Helsinki and was approved by the local ethics committee of the University of Leipzig. Prior to the study, potential participants completed a screening interview by telephone. Inclusion criteria were female gender and age of 20-35 years old. Exclusion criteria comprised a history of neuropsychiatric disorders –such as depression (Beck Depression Inventory, BDI > 18) – medical disorders such as hypertension, hyper- or hypothyroidism, cancer, and diabetes, as well as left-handedness and MRI incompatibilities.

During the recruitment process, two additional participants had to be excluded due to pathologies not detected during the screening interview, one additional participant declined to perform the MRI session, and six additional participants were removed from the fMRI analysis due to excessive head movements (framewise displacement > 0.2). All participants gave written informed consent prior to taking part in the study and received a compensation for their time in the study.

### Anthropometric measures

On the day of the fMRI acquisition, we measured participants’ weight, height, waist circumference and hip circumference. We used waist circumference as a surrogate measure of adiposity in our analyses. Nevertheless, we repeated all the analyses using BMI instead of waist circumference and the results did not change (data not shown).

### Self-report questionnaires

A week before the date of the MRI acquisition, participants received an internet-based survey (using the Limesurvey platform, www.limesurvey.org). The data collection for the present project was coordinated with a second independent project aimed at tackling social processing in obesity, whose results will be presented elsewhere. Participants completed several questionnaires that addressed social support and withdrawal as well as personality. We also administered the Three Factor Eating Questionnaire (TFEQ), which provides three measurements of eating behavior: cognitive restraint over eating, disinhibited eating and hunger (Stunkard and Messick, 1985). We evaluated chronic stress with the Trierer Inventar zum Chronischen Stress (TICS) (Schulz et al., 2004). The total time estimated for the completion of the Internet survey was 45 minutes.

### Emotional priming task

Participants completed an fMRI event-related food processing task, in which we presented food and object photographs preceded by priming emotional stimuli. The task consisted of 6 conditions, combining 3 types of priming (negative, neutral, and positive) and 2 types of stimuli (food and objects). Participants viewed an emotional picture (negative, neutral or positive) for 4 seconds. After a 2 seconds inter-stimuli interval with a blank screen, we presented either a food photograph or an object photograph for another 2 seconds. Subsequently, participants were asked to rate how much they liked the food/object on a −5 to +5 Likert scale, by utilizing a button box placed in their right hand. The Likert scale was programmed to last 8 seconds maximum, but as soon as the participants performed the rating, a fixation cross appeared on the screen for the remainder of the 8 seconds (Figure 1). Each condition consisted of 15 trials, and the order of the conditions was semi-randomized across participants – i.e., to ensure that the same condition would not appear more than twice for every 10 trials.

**Figure 1:**
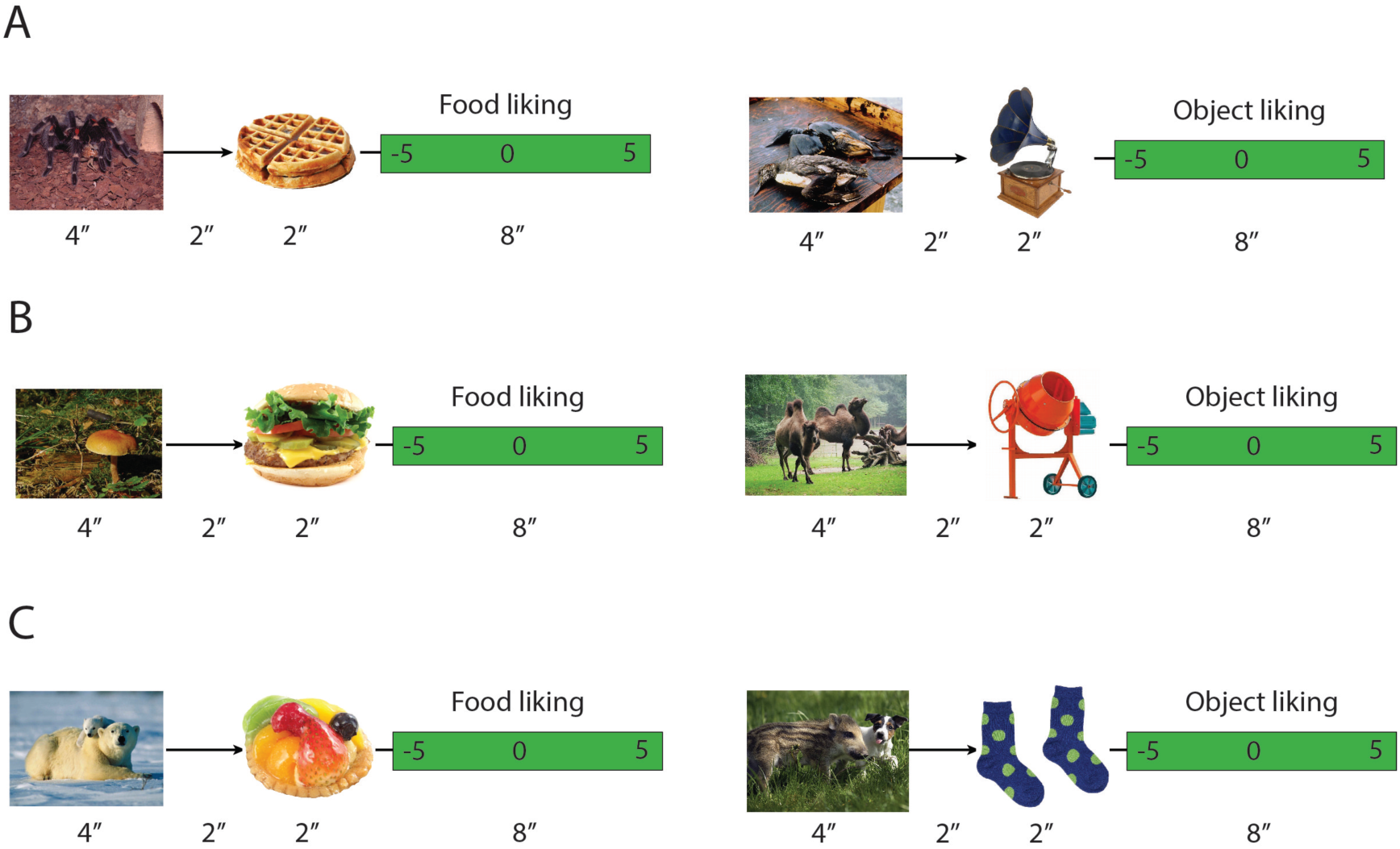
Schema of the fMRI task. We presented pictures of foods and objects preceded by emotional priming images. A) Negative emotional priming. B) Neutral emotional priming. C) Positive emotional priming.

Before the task we collected hunger rates (rated from 1 to 10, with higher values representing higher hunger) and number of hours before their last meal.

The emotional stimuli were extracted from the Emotional Picture Set (Wessa, M., Kanske, P., Neumeister, P., Bode, K., Heissler, J., & Schönfelder, 2010), a dataset that provides normative data from emotional images rated by German population in terms of valence and arousal. We selected the images rated as lowest, neutral, and highest in terms of valence for the negative, neutral, and positive emotional categories respectively. Repeated measures ANOVAS showed that the three image categories were similar in terms of luminosity, contrast, and color saturation (lowest Bonferroni corrected p = 0.9). Similar to the population ratings, the valence and arousal of the images was significantly different between the three groups (all Bonferroni corrected p<0.001).

We selected food and object photographs from the FoodCast Research Image Database (FRIDa) (Foroni et al., 2013). Food and objects were not significantly different in spatial frequency, brightness or size (lowest p = 0.065 after Bonferroni correction). However, and similar to the population ratings, valence and arousal of food items were higher than of objects (Bonferroni corrected p=0.005 for both tests). After the MRI session, participants rated each image regarding valence and arousal using a 1-9 Likert scale.

### MRI acquisition

Images were acquired on a 3 Tesla Skyra scanner (Siemens, Erlangen, Germany). Eight hundred seventeen whole brain fMRI volumes were acquired using a multi-slice gradient-echo EPI sequence during 27.20 minutes [echo time (TE): 22 ms; repetition time (TR): 2000 ms; 2.5 mm slice thickness; 40 slices per volume; 20% interslice gap; 90° flip angle; 192 mm field of view (FOV); voxel size 3.0 × 3.0 × 2.5 mm^3^].

For each participant, we also acquired a T1-weighted anatomical image with an MPRAGE ADNI protocol with 9.14 minutes of duration (T1: 900 ms; TE: 2.01 ms.; TR: 2300 ms.; 1-mm slice thickness; 50% interslice gap; 9° flip angle; 256 mm FOV; bandwidth: 240 Hz/pixel; sagittal orientation; voxel size: 1.0 × 1.0 × 1.0 mm^3^).

### FMRI analysis

We preprocessed and analyzed MRI data using SPM12 (Wellcome Trust Centre for Neuroimaging, UCL, London, UK), implemented in Matlab (The MathWorks Inc., Sherborn, MA). First, fMRI volumes were unwarped and spatially aligned to the first image of the session. Realignment parameters (motion parameters) were calculated and later used as nuisance regressors. We then performed slice-timing correction to the anatomical middle slice and coregistered the fMRI data to the high-resolution anatomical image. The reference image was segmented into gray matter, white matter and cerebrospinal fluid, and the MRI images were normalized to Montreal Neurological Institute (MNI) standard space. Finally, we applied an isotropic Gaussian kernel of 8 mm FWHM to smooth the normalized images.

For each participant, we modeled on the first level (a) the onset of each category of emotional stimuli (b) the onset of food and objects. That is, our design had 3 regressors modeling priming stimuli (negative, neutral and positive) and 6 regressors modeling the events of interest: (i) food presented after negative priming; (ii) objects presented after negative priming; (iii) food after neutral priming; (iv) objects after neutral priming; (v) food after positive priming; (vi) objects after positive priming. All six-motion parameters were modeled as nuisance regressors. Fixation trials and the time corresponding to perform the liking rates for foods/objects were left unmodeled.

First, we tested if priming stimuli elicited brain activity in coherence with what has been previously described in the literature. For that, we included the contrasts “negative emotional images > neutral images” and “positive emotional images > neutral images”. We did not include the comparison between negative images and positive images since this contrast does not usually yield a robust pattern of brain activity (Lindquist et al., 2015). Our main contrast of interest was the difference in fMRI activity between foods and objects. This contrast is widely used in neuroimaging studies of nutrition to examine brain activity that is specific to food (van der Laan et al., 2011). We assessed this contrast (i.e., foods > objects) for each category of emotional priming (e.g., negative priming: foods > objects).

In a second-level analysis, we performed one-sample t-tests to examine the effect of each contrast. In an exploratory analysis, we included interaction contrasts to test for the whole-brain differences between foods and objects across priming conditions. For all these contrasts, we tested for the effects of obesity by including waist circumference as a regressor of interest. Since the participants reported a heterogeneous number of hours fasting, we included subjective rates of hunger as a regressor. We also added mean framewise displacement, calculated from the movement parameters (Power et al., 2012), as a nuisance regressor. Statistical threshold was set at FWE corrected p<0.05 at the voxel level.

### Amygdala analysis

We tested whether amygdala activity for the contrast “food stimuli > objects” was influenced by emotional priming stimuli. We used the MarsBaR (MARSeille Boîte À Région d’Intérêt) toolbox from SPM to extract data from the amygdala. The amygdala ROI was defined with the Harvard-Oxford Atlas. In brief, MarsBaR allows calculating the mean for all the voxels included in the ROI. This produces time-course values for each subject and for each regressor specified in the first level analysis. We extracted the time courses associated with our events of interest (6 events that are the combination of 3 emotional priming (negative, neutral, positive) and the 2 types of stimuli (food, objects)). With these values, we run the statistical model in R. Since we were primarily interested in the contrast “food stimuli > objects”, we subtracted the values of foods and objects in the amygdala separately for each emotional priming condition.

### Statistical analysis: liking rates and amygdala activity

We tested whether emotional priming modifies liking rates and amygdala activity in response to foods compared to objects. We performed a repeated measures design with a multilevel linear model and compared three conditions (i.e., (1) negative priming: foods – objects; (2) neutral priming: foods – objects; (3) positive priming: foods – objects) using the R function lme() (package nlme). We performed post-hoc tests by applying Tukey’s correction. We performed post-hoc orthogonal contrasts to examine the difference between foods and objects during the negative versus neutral priming and positive versus neutral priming.

## Results

### Demographical characteristics

We included fifty-eight female participants in the analyses; see Table 1 for a distribution of the demographic and clinical variables. Among them, 2 participants were underweight (BMI < 18.5), 30 participants were normal-weight (18.5 ≤ BMI < 25), 13 participants were overweight (25 ≤ BMI < 30) and 12 participants were obese (BMI ≥ 30).

**Table 1.** Demographic characteristics and psychological measures.

| Table 1. Demographic characteristics and psychological measures. |  |  |  |  |
| --- | --- | --- | --- | --- |
|  | Minimum | Maximum | Mean | Std.<br>Deviation |
| Age | 20.0 | 35.0 | 26.33 | 3.69 |
| BMI ($\text{kg/m}^2$) | 17.67 | 46.83 | 25.63 | 5.84 |
| Waist circumference (cm) | 61 | 104 | 76.57 | 10.44 |
| Waist-to-hip ratio | .69 | .93 | .81 | .05 |
| Subjective hunger (1-10) | 1 | 7 | 2.1 | 1.66 |
| Hours fasting | 0.1 | 12 | 2.16 | 2.45 |
| BDI | .0 | 12.0 | 4.05 | 3.48 |
| Chronic stress (TICS) | 13 | 162 | 81.72 | 28.40 |
| Cognitive restraint (TFEQ) | 0 | 17 | 6.88 | 4.52 |
| Disinhibited eating (TFEQ) | 1 | 14 | 6.21 | 3.40 |
| Hunger (TFEQ) | 0 | 13 | 4.07 | 3.14 |

### Post-scan ratings for valence and arousal

In line with the normative ratings, the emotional images (negative, neutral and positive) used for priming differed both in valence (X^2^(2)=403.19; p<0.001) and arousal (X^2^(2)=156.11; p<0.001) ratings in our sample. Likewise, foods and objects differed both in valence (X^2^(1)=8.91; p<0.003) and arousal (X^2^(1)=41.80; p<0.001) ratings, with food stimuli obtaining higher scores for both measures (Figure 2).

**Figure 2.**
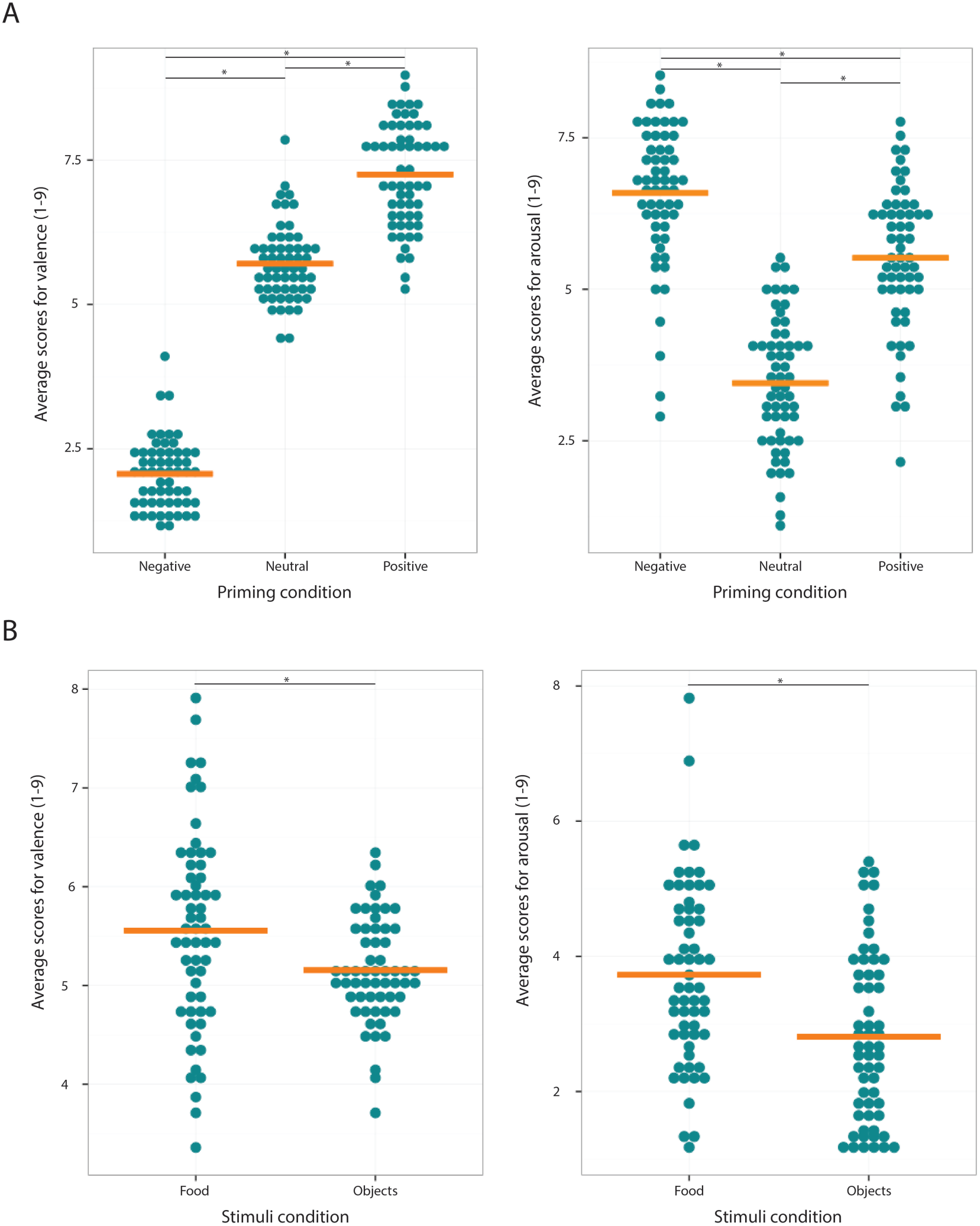
Average scores for valence and arousal provided after the MRI scan. (A) Scores for each priming emotional stimuli (negative, neutral and positive). (B) Average scores for food and object stimuli. Asterisks depict significant post-hoc differences.

### Liking rates

The multilevel model for repeated measures showed that priming condition had an effect on the difference between liking scores for food and objects X^2^(2)=6.286; p=0.0432. Orthogonal contrasts showed that the difference between liking scores for foods and objects was higher during the positive priming condition compared to the neutral condition (b=0.184< t_(114)_=2.41; p=0.018). The contrast between the negative and the neutral priming was non-significant (b=0.140; t_(114)_=1.84; p=0.068) (Figure 3). Participants’ waist circumference had no significant effect on the liking rates, X^2^(3)=0.01; p=0.945.

**Figure 3.**
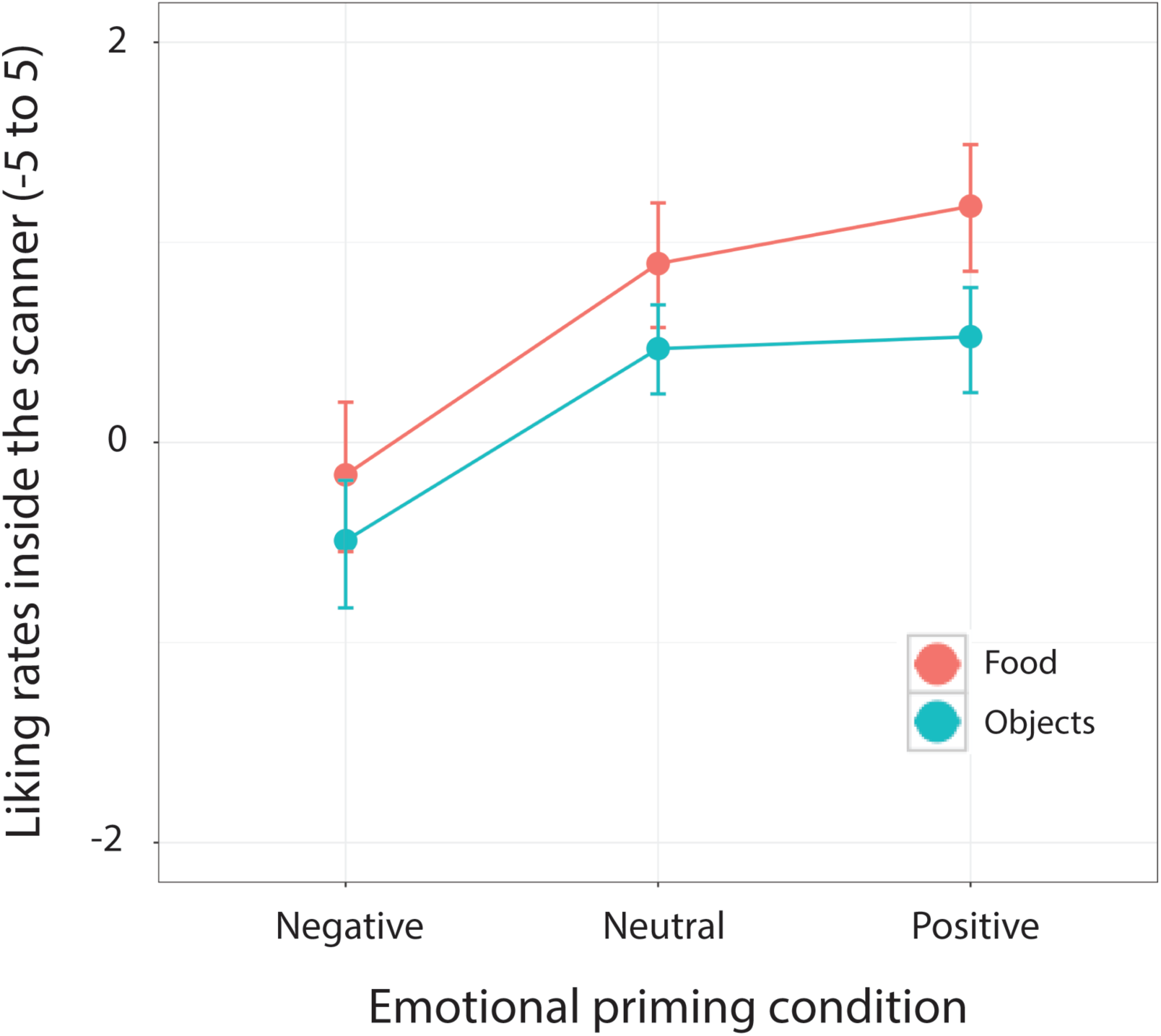
Liking rates for food and objects during the negative, neutral and positive priming condition. The error bars represent 95% confidence intervals.

### Neuroimaging results

The emotional priming task contained two sets of images: emotional images (priming stimuli) and the events of interest (foods and objects under each emotional priming condition).

#### 0. Effects of emotional images

In order to test that the emotional images were able to yield a pattern of activity in line with what it has been described in the literature, we first examined the contrasts “negative emotional stimuli > neutral stimuli” and “positive emotional stimuli > neutral stimuli”. Relative to neutral stimuli, negative emotional images activated the left amygdala, bilateral temporal occipital fusiform, brainstem right precentral gyrus, right inferior frontal gyrus, left OFC and left lateral occipital cortex. The contrast positive > neutral stimuli activated the bilateral lateral occipital cortex and the precuneus (Table 2).

**Table 2.** Whole brain results for the emotional priming stimuli

| Table 2. Whole brain results for the emotional priming stimuli |  |  |  |  |  |
| --- | --- | --- | --- | --- | --- |
| Brain region | MNI Coordinates |  |  | Cluster size | T |
|  | X | Y | Z |  |  |
| *Negative emotional stimuli > Neutral stimuli* |  |  |  |  |  |
| Left amygdala | -20 | -6 | -12 | 654 | 9.90 |
| Right temporal occipital fusiform | 46 | -42 | -18 | 1027 | 9.44 |
| Left temporal occipital fusiform | -40 | -50 | -16 | 734 | 9.18 |
| Brain stem | -6 | -28 | -10 | 370 | 8.76 |
| Right precentral gyrus | 44 | 8 | 28 | 239 | 8.09 |
| Right inferior frontal gyrus | 50 | 34 | 8 | 244 | 7.68 |
| Left OFC | -28 | 12 | -18 | 61 | 7.11 |
| Left lateral occipital cortex | -46 | -64 | 16 | 107 | 6.79 |
| *Positive emotional stimuli > Neutral stimuli* |  |  |  |  |  |
| Right lateral occipital cortex | 44 | -80 | -10 | 1147 | 11.01 |
| Left lateral occipital cortex | -38 | -86 | -10 | 913 | 10.38 |
| Precuneus | 0 | -60 | 32 | 63 | 5.98 |

Results are FWE corrected at the voxel level $p < 0.05$ ; minimum cluster size: 50 voxels.

#### 1. The contrast foods > objects: differences depending on the emotional priming condition

Under negative priming, the contrast foods > objects yielded activity in the occipital pole and lateral orbitofrontal cortex (OFC) bilaterally. After the presentation of neutral images, we found that the contrast foods > objects engaged bilateral activity in the occipital pole, lateral occipital cortex, insula and amygdala. Finally, during the positive priming condition, the contrast foods > objects was associated with bilateral activity in the lateral OFC, occipital pole, insula, as well as with left lateralized activity in the amygdala (Table 3; Figure 4). In an exploratory whole-brain analysis, we did not find significant differences between the priming conditions in the contrast “foods > objects”. Neither waist circumference nor hunger rates had a significant effect in any of the contrasts.

**Figure 4.**
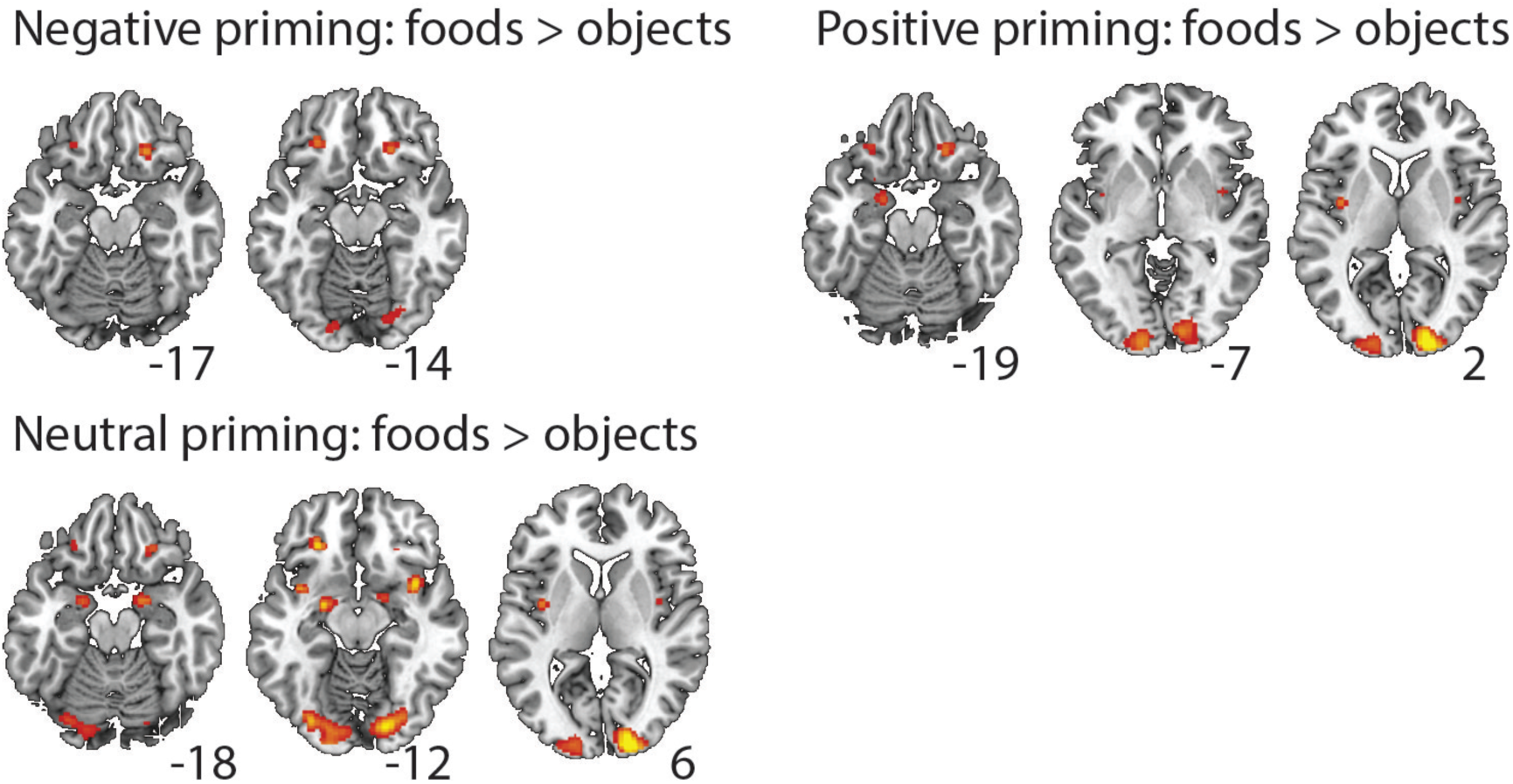
Whole brain fMRI results for the contrast “foods > objects” during the different priming conditions.

**Table 3.** Whole brain results for the events of interest

| Table 3. Whole brain results for the events of interest |  |  |  |  |  |
| --- | --- | --- | --- | --- | --- |
| Brain region | MNI Coordinates |  |  | Cluster size | T |
|  | X | Y | Z |  |  |
| *Negative priming condition: Foods > Objects* |  |  |  |  |  |
| L occipital pole | -12 | -94 | -2 | 588 | 9.18 |
| R occipital pole | 12 | -90 | -4 | 581 | 8.44 |
| R lateral OFC | 20 | 30 | -16 | 83 | 8.31 |
| L lateral OFC | -26 | 34 | -14 | 96 | 7.92 |
| *Neutral priming condition: Foods > Objects* |  |  |  |  |  |
| R occipital pole | 14 | -92 | 0 | 2414 | 11.82 |
| R insula | 38 | 8 | -12 | 148 | 8.71 |
| L lateral OFC | -26 | 34 | -14 | 94 | 8.69 |
| L amygdala | -22 | -6 | -12 | 245 | 8.13 |
| R amygdala | 20 | -2 | -16 | 113 | 8.13 |
| L insula | -36 | -4 | 10 | 90 | 7.86 |
| R lateral OFC | 24 | 30 | -16 | 53 | 7.20 |
| *Positive priming condition: Foods > Objects* |  |  |  |  |  |
| R lateral OFC | 22 | 28 | -16 | 132 | 10.05 |
| L lateral OFC | -26 | 32 | -16 | 130 | 9.53 |
| L occipital pole | -12 | -94 | -8 | 694 | 9.33 |
| R insula | 38 | 4 | -12 | 132 | 8.21 |
| R occipital pole | 16 | -92 | 2 | 663 | 8.20 |
| L insula | -38 | -6 | 10 | 67 | 7.32 |
| L amygdala | -18 | -4 | -22 | 79 | 7.30 |
| Results are FWE corrected at the voxel level $p < 0.05$; minimum cluster size: 50 voxels. | | | | | |

#### 2. Interactions in the amygdala

Emotional priming had a significant effect on the difference between fMRI activity in the left amygdala in response to food stimuli and objects, X^2^(2)=6.374; p=0.041. Orthogonal contrasts showed that differences in fMRI activity between foods and objects were significantly higher during neutral priming compared to negative priming, b=-0.023; t_(114)_=-2.313; p=0.023. There was no difference between positive and neutral priming (b=-0.003; t_(114)_=-0.268; p=0.789) (Figure 5). There was no effect of priming in the right amygdala.

**Figure 5.**
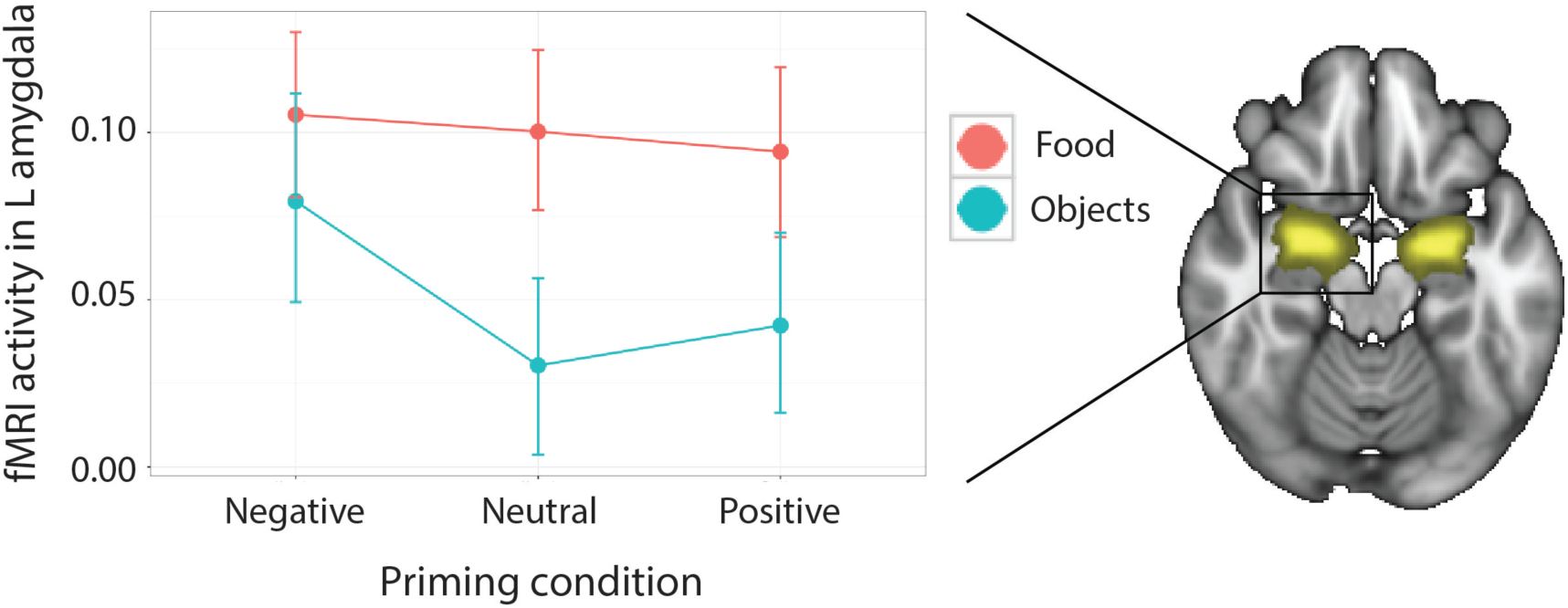
Emotional priming shows a significant effect on the difference between fMRI activity in the left amygdala in response to foods and objects.

Waist circumference had an additional effect on fMRI activity in the left amygdala X^2^(4)=4.754; p=0.029. A closer examination into this effect indicated that, during the neutral priming condition, waist circumference was inversely related to the difference in left amygdala activity between food stimuli and objects (r=-0.40; Bonferroni corrected p=0.006). There was no effect of waist circumference in the other conditions. Hunger rates did not have an effect on amygdala activity.

Finally, we performed exploratory correlations between amygdala activity and a set of other potential variables of interest (i.e., liking scores, cognitive restrain over eating (TFEQ), stress (TICS) and depression (BDI)). These correlations were performed with the sole intention of guiding future research on the topic. After correcting for multiple comparisons, all the correlations were not significant. There was a non-significant trend in the correlation between stress scores and amygdala activity during the positive emotional condition (r=0.30, uncorrected p=0.02).

## Discussion

The current study examined whether emotional context can influence the neural processing of food stimuli. To this aim, we designed an fMRI task, in which emotional priming stimuli (pictures of negative, neutral and positive situations) were followed by images of foods and objects. We obtained subjective liking scores for foods and objects and examined the contrast “foods > objects” across the different emotional contexts. We found that there was an overall effect of priming on the difference between foods and objects in terms of liking scores. Moreover, this difference was higher after the positive priming condition than after the neutral priming condition. In an exploratory whole-brain analysis, we did not find significant effects of emotional context on food processing. Since the amygdala was our main ROI, we performed further analyses by extracting fMRI signal changes in this area. We found a significant effect of priming in the left amygdala activity in the difference between foods and objects. This difference was higher after the neutral emotional prime than after the negative emotional prime. Waist circumference, however, did not have an effect on emotional food processing. Our results suggest that emotional context modifies the salience advantage of food, given the role of the amygdala in salience detection. Food seems to have a certain salience advantage over objects under positive and neutral circumstances. However, under negative circumstances, foods and objects might be assigned a similar amount of attentional resources.

### Neurobehavioral effects of emotional context on food processing

Eating can be a potent pleasant and rewarding behavior. The rewarding value of food arises from signals in the mesolimbic dopamine circuit, a brain system that drives approach responses towards food and other rewards (Haber and Knutson, 2010). The processing of food, however, is not rigid but determined by internal and external factors of the context. Hunger, for instance, is a factor that can impact on the way that mesocorticolimbic areas process food (Charbonnier et al., 2018). In the current study, we have focused on emotional context, a variable that had been shown to impact eating behaviors. Participants in our study rated foods higher than objects in terms of liking. This difference between food and objects, however, was bigger after positive priming compared to neutral priming. It also showed a trend for higher liking rates after neutral compared to negative priming. Our findings thus suggest that emotional context modifies the advantage of foods over objects in terms of liking.

Emotions, reward and pleasure are intricately connected. For instance, anhedonia –i.e., decreases in the ability to experience pleasure – might occur as a consequence of negative emotional states. Using a threat-of-shock task, a behavioral study in healthy women showed that acute stress impaired responsiveness on a monetary decision-making task (Bodgan and Pizzagalli, 2016). Another behavioral study found that, when participants listened to music that they disliked, they rated chocolate ice cream as being less pleasant. When the same participants listened to music that they liked, they rated chocolate ice cream as being more pleasant (Kantono et al., 2016). Across all the emotional priming conditions, the contrast “foods > objects” yielded a pattern of brain activity that was closely in line with previous findings. This pattern included the occipital cortex, lateral OFC, insula and amygdala. Similar to our findings, a meta-analysis on 17 studies examined the neural correlates of food pictures and reported a consistent engagement of the posterior fusiform gyrus, lateral OFC and the insula. The same paper found that the amygdala was additionally engaged when participants were hungry (van der Laan et al., 2011). This possibly reflects increases in the salience processing (or behavioral relevance) of food during hunger. The amygdala is a brain region highly involved in the processing of emotional stimuli as well as in food processing. For this reason, we chose the amygdala as our main ROI. In the left amygdala, the difference in fMRI activity between foods and objects changed according to the emotional priming condition. Food, relative to non-food processing, was associated with a greater fMRI activity in the neutral and positive priming conditions, with no significant differences across these two conditions. The advantage that food presented over objects, however, was reduced during the negative emotional priming. In this condition, foods and objects showed a similar level of activity in the left amygdala.

Detecting and processing negative emotional stimuli in an efficient manner is crucial for well-being and survival. One possible interpretation for our findings in the amygdala is that a negative emotional condition might decrease the salience advantage of food. Hypothetically speaking, when an organism perceives negative emotional stimuli, it might elicit a general state of alertness to provide appropriate responses to the stimuli (such as fight or flight). This general state of alertness might overshadow some important stimuli, like food, that become comparatively less crucial. This way, foods and objects might be assigned a similar amount of salience resources – which is also suggested by comparable amounts of activity in the left amygdala. Conversely, when the emotional context is neutral or positive, the organism might be in a more favorable position to discriminate between the processing of rewarding stimuli (e.g., food) and neutral stimuli such as objects. In this vein, acute stressors seem to suppress appetite responses by engaging the hypothalamic-pituitary-adrenal axis (Sominsky and Spencer, 2014). An interesting possibility here is that the similar processes might apply when replacing the food stimuli with another motivating stimuli, such as monetary reward. While we could not find direct evidence supporting this hypothesis, a previous study suggests that emotional cues enhances monetary loss aversion responses in the amygdala and striatum in individuals reporting low levels of anxiety (Charpentier et al., 2016).

Previous studies have applied stress paradigms (a condition associated with negative emotionality) to evaluate the role of the amygdala and other corticolimbic areas for food-related behaviors. In this vein, two studies have suggested that, under stress, the amygdala and other mesocorticolimbic areas are *less engaged* in response to food. The first study, conducted in healthy female participants, suggested that, relative to a neutral condition, participants in a stressful condition exhibited lower activation in amygdala, cingulate cortex, and hippocampus during a food choice task (Born et al., 2009). A second study, which examined women with bulimia nervosa symptoms, showed that being in an experimental stressful situation was associated with decreases in BOLD signal in the anterior cingulate cortex, amygdala and ventromedial prefrontal cortex in response to food cues, as compared to a non-stressful control condition (Fischer et al., 2017). Moreover, these decreases in BOLD signal seemed to act as mediators on the relationship between increased negative affect and binge eating episodes (Wonderlich et al., 2018). However, evidence for the opposite effect, and findings that activity in the amygdala is *higher* during stress, also exist. Using an fMRI food choice paradigm, a study on male participants reported that, compared with participants in a neutral condition, subjects assigned to a stressful condition put greater weight on the taste of the food items presented. In parallel, bilateral amygdala and right nucleus accumbens reflected the relative taste value of chosen options more strongly in stressed compared to control participants. The authors interpreted these findings as suggesting that stress may increase the effect of the rewarding attributes of the stimuli (Maier et al., 2015). Finally, another study examined fMRI activity in response to food stimuli in university students (both sexes) during the final exams and during a non-exam period. Personality differences in stress reactivity, as measured by the behavioral inhibition scale, predicted increases in perceived stress during the final exams. Moreover, higher scores in the behavioral inhibition scale were associated with increased fMRI activity during the exam versus the non-exam period in the amygdala and ventromedial prefrontal cortex in response to foods (Neseliler et al., 2017).

Methodological heterogeneities between studies limit the comparability between the current studies and previous fMRI studies in stress. The most obvious one is that stress paradigms and emotional priming paradigms might not produce equivalent emotional reactivity. Sex is another factor that limits the generalizability of our results, since we only recruited women in our study.

### Weight status does not have an effect on the interaction between emotional context and food processing

Overweight and obesity may play an important role in the interplay between emotions and food consumption. Previous studies have reported that participants’ body mass affects their food intake and weight gain during chronic stress (Fujishiro et al., 2015; Kivimäki et al., 2006). In our statistical models, however, waist circumference did not have an impact on liking scores. In the case of the amygdala analysis, the effect of waist circumference was driven by the neutral (non-emotional) condition. Our findings thus do not support the hypothesis that waist circumference has a differential influence on food processing according to the emotional context. However, more studies are needed before this statement is conclusive.

The results of our study should be read with caution in view of several limitations. First, we did not recruit male participants, which limits the generalizability of our results. Second, the sample size is relatively small to evaluate the effects of other factors that might have an effect on emotional processing and food, such as cognitive control over eating behavior, perceived stress, depressive symptoms or menstrual cycle. Moreover, the current sample size might not be adequate to detect whole-brain differences between emotional priming conditions in the contrast between food and non-food stimuli. Third, we did not evaluate the possible presence of clinical or subclinical eating disorders, which might have had an effect on food processing. We hope that future studies will address these issues.

### Conclusions

In the present study we have addressed two main questions: (i) how does emotional context (negative, neutral and positive) influence the neurobehavioral processing of food and non-food stimuli; and (ii) whether waist circumference (a surrogate measure of abdominal fat) affects the interplay between emotional stimuli and food processing. Our findings suggest that liking scores change with the valence of the emotional context. Liking scores for both food and non-food stimuli were lowest after negative priming and increased after neutral and positive priming, with overall higher scores for food compared to non-food stimuli. Most importantly, the difference in liking scores between foods and objects was the highest during the positive emotional context. That is, positive priming had a stronger effect on food compared to non-food stimuli. Similar to this, the difference in amygdala activity between foods and objects was lower during the negative emotional context than during the neutral emotional context. This is the first time that the interactions between emotional priming and food processing have been observed both behaviorally and with fMRI. Our findings emphasize the importance of emotional experiences in the context of food processing, and might inform clinical research and interventions. Future lines of research could extend these findings to patients with eating disorders.

## Acknowledgements

We thank all the participants for their collaboration in the study. We also thank Bettina Johst, Ramona Menger and Nicole Pampus for technical support during the preparation of the study and data recruitment.

IGG is supported by a Postdoctoral Fellowship from the Canadian Institutes of Health Research. MG is supported by the Federal Ministry of Education and Research (grant 13GW0206B). AD is funded by the Canadian Institutes of Health Research. AH is supported by the Deutsche Forschungsgemeinschaft (DFG, German Research Foundation) – project number 209933838 – SFB 1052 and by the German Federal Ministry of Education and Research (FKZ: 01EO1501). JN was supported by the German Federal Ministry of Education and Research (FKZ: 01EO1001). None of the authors declares a potential conflict of interest.

## Open practice statement

The study was not formally preregistered. The analysis code and task-related materials are available in Open Science Framework (https://osf.io/v49ez/). Whole-brain unthresholded T maps are available in NeuroVault Nutritional collection (https://identifiers.org/neurovault.collection:5964). The dataset will be available from the corresponding author on reasonable request.

## Notes

https://osf.io/v49ez/

https://identifiers.org/neurovault.collection:5964

